# MechFind: A computational framework for de novo prediction of enzyme mechanisms

**DOI:** 10.1101/2025.09.25.678649

**Authors:** Austin D Hartley, Vikas Upadhyay, Veda Sheersh Boorla, Costas D Maranas

## Abstract

Despite the importance of understanding the step-by-step mechanism of enzymatically catalyzed reactions, fewer than one thousand cataloged mechanistic annotations can be found in the open literature and databases. Herein, we introduce MechFind, a computational tool that generates elementally and charge-balanced putative enzyme mechanisms. Unlike previous methods that require structural data or user-supplied active site residues, MechFind uses only the overall reaction stoichiometry as input, abstracting individual reaction steps as the gain or loss of chemical moieties. An optimization framework then identifies the top ten most parsimonious (i.e., fewest steps) mechanistic descriptions for the overall transformation, which are then re-ranked based on their mechanistic similarity to a database of known, curated mechanisms. MechFind recovered the correct mechanism for 72% of the Mechanism and Catalytic Site Atlas (M-CSA) training dataset within the top ten predictions and was independently validated on six enzyme mechanisms absent from the training set. When deployed at scale on 28,412 reactions from the Rhea database, MechFind identified a plausible mechanism for 64% of all entries, generating over 18,000 novel mechanistic hypotheses expanding significantly upon the current state of the art in prediction (i.e., a 20-fold increase). By proposing detailed reaction mechanisms MechFind provides information that can be leveraged by modern ML-based de novo protein design tools, offering a resource for improving functional annotation and accelerating the engineering of novel biocatalysts. All codes, curated datasets, and results are available at https://github.com/maranasgroup/MechFind.

## Main Text Introduction

Enzymes catalyze nearly every biochemical transformation in living systems, mediating both the catabolic oxidation of carbon substrates to harvest energy and the anabolic assembly of small molecules into the macromolecular machinery of the cell. Their remarkable chemoselectivity and rate enhancements have been harnessed in biotechnology to produce a broad array of compounds, from bulk biochemicals such as amino acids [1], vitamins [2] and polymer monomers [3] to high-value pharmaceuticals [4], vaccine adjuvants [5] and pheromones [6].The ability to engineer existing enzymes or design wholly new biocatalysts is therefore of significant value, but has traditionally depended on time- and cost-intensive methods such as random mutagenesis [7] and directed evolution [8]. However, recent breakthroughs in de novo protein modeling [9] [10], backbone generation [11], and sequence design [12] have begun to accelerate progress towards effective enzyme redesigns. Successful examples of computationally driven enzyme design include functional luciferases [13], Kemp eliminases [14], and serine hydrolases [15], with many more highlighted in review articles [16] [17] [18]. These examples use knowledge of the reaction’s transition state to design possible “theozymes” which are computational models of the active site. According to transition state theory [19] [20], an enzyme achieves its catalytic power by preferentially stabilizing this transition state relative to the substrate and product complexes. Knowledge of the transition state can, in turn, be computed from the enzyme’s mechanism, as each mechanistic step describes the flow of electrons and the rearrangement of atoms and bonds.

Therefore, a complete, stepwise understanding of enzyme mechanisms is essential for illuminating the transition states needed to engineer new active sites for novel reactions. Yet, a significant “mechanism gap” exists as fewer than one thousand enzymatically catalyzed reaction entries collected in major biochemical repositories carry complete mechanistic annotation. For example, the Mechanism and Catalytic Site Atlas (M-CSA) [21] curates thousands of examples but only 734 entries document every bond-breaking and bond-forming event in a complete mechanism. This scarcity stands in stark contrast to resources such as USPTO [22], BRENDA [23], KEGG [24], Rhea [25] or MetaNetX [26], which catalog tens of thousands of balanced reactions, though without any mechanistic detail. Bridging this “mechanism gap” is a significant obstacle to expanding enzyme design beyond a few model systems. Prior computational efforts, such as MechSearch [27] and EzMechanism [28], partially address this challenge but are limited by their requirement for user-supplied active-site residues and, in the case of EzMechanism, a high-resolution protein structure. MechSearch has been used to propose mechanisms for less than a thousand reactions from the Rhea database. Moreover, their graph-based representations of elementary steps can become computationally prohibitive when mined exhaustively across millions of hypothetical enzyme-substrate pairs.

Herein, we introduce MechFind, a computational framework that predicts detailed enzymatic mechanisms using only the chemical structures of the starting materials and final products. At its core lies a moiety-based encoding [29] in which every non-hydrogen atom is classified by its local bonding environment, and each reaction is summarized as a vector of “moiety gains” and “moiety losses.” This approach was inspired by the novoStoic [30] framework, which used a similar encoding to design multi-step metabolic pathways; MechFind adapts this concept to predict multi-step enzyme mechanisms. The original optimization formulation carries out both the identification and ordering of the elementary steps. However, we found that it is more computationally efficient to split these two tasks into two separate optimization tasks. To distinguish the correct mechanism from other chemically plausible, parsimonious alternatives, we also developed a re-ranking procedure that scores each candidate based on its similarity to known mechanisms. This method identifies the most plausible candidate by finding the one that most closely resembles a validated mechanism from our curated database. Through this hybrid approach, MechFind recovered the correct mechanism among the top ten predictions for 85% of the 661 entries in the M-CSA [21] database deemed valid inputs. MechFind was independently validated on a set of six recently published mechanisms absent from the training set [31–36]. When deployed at scale on 28,412 reactions from the Rhea database [25], MechFind identified a plausible mechanism for 64% of all entries, generating over 18,000 novel mechanistic hypotheses and systematically characterizing failures as either limitations in computational tractability or gaps in the underlying moiety description set. Finally, we demonstrate MechFind’s utility not just as a prediction tool but as an exploratory engine to map the network of competing catalytic strategies for a given reaction, providing a rich set of testable hypotheses for de novo enzyme design.

## Results

We begin by benchmarking the performance of the initial parsimony-based approach on the curated M-CSA database [21]. The parsimony-based framework, on its own, successfully recovered the correct mechanism as the top-ranked prediction in 64% of cases and within the top ten predictions in 85% of cases. We then test the framework’s capacity for discovery by validating it against a set of six recently characterized mechanisms absent from the training data, successfully identifying the correct mechanism in all cases. Having assessed its accuracy, we then deploy MechFind at a large scale to annotate tens of thousands of reactions from the Rhea [25] and MetaNetX [26] databases, generating over 18,000 novel mechanistic hypotheses and representing a more than 20-fold increase in available mechanisms. Finally, we contextualize our results by comparing MechFind’s performance directly with existing tools and demonstrate its utility as an exploratory engine for mapping the landscape of mechanistic diversity for a given reaction.

### Moiety-Based Representation of Reactions

Each M-CSA [21] entry enumerates substrates, products, catalytic residues/cofactors, and formal arrow-pushing schemes for every step. MechFind abstracts this information into a moiety-based representation. For simplicity, we represent each moiety using a canonical SMILES string [37]. Every elementary rule is defined by its changes in moiety counts from the set of all unique moieties M. Each unique rule is placed in a matrix denoted, where entry represents the gain (positive value) or loss (negative value) of moiety m according to rule r. The overall reaction stoichiometry is similarly encoded as a vector which describes the net moiety changes between all products and reactants. The fundamental requirement of any valid mechanism is that the cumulative action of all its elementary steps must match the overall reaction stoichiometry. Figure 1 illustrates this encoding process, where the sum of the moiety change vectors for the elementary steps (Figure 1b) equals the moiety change vector for the overall reaction (Figure 1a).

**Figure 1.**
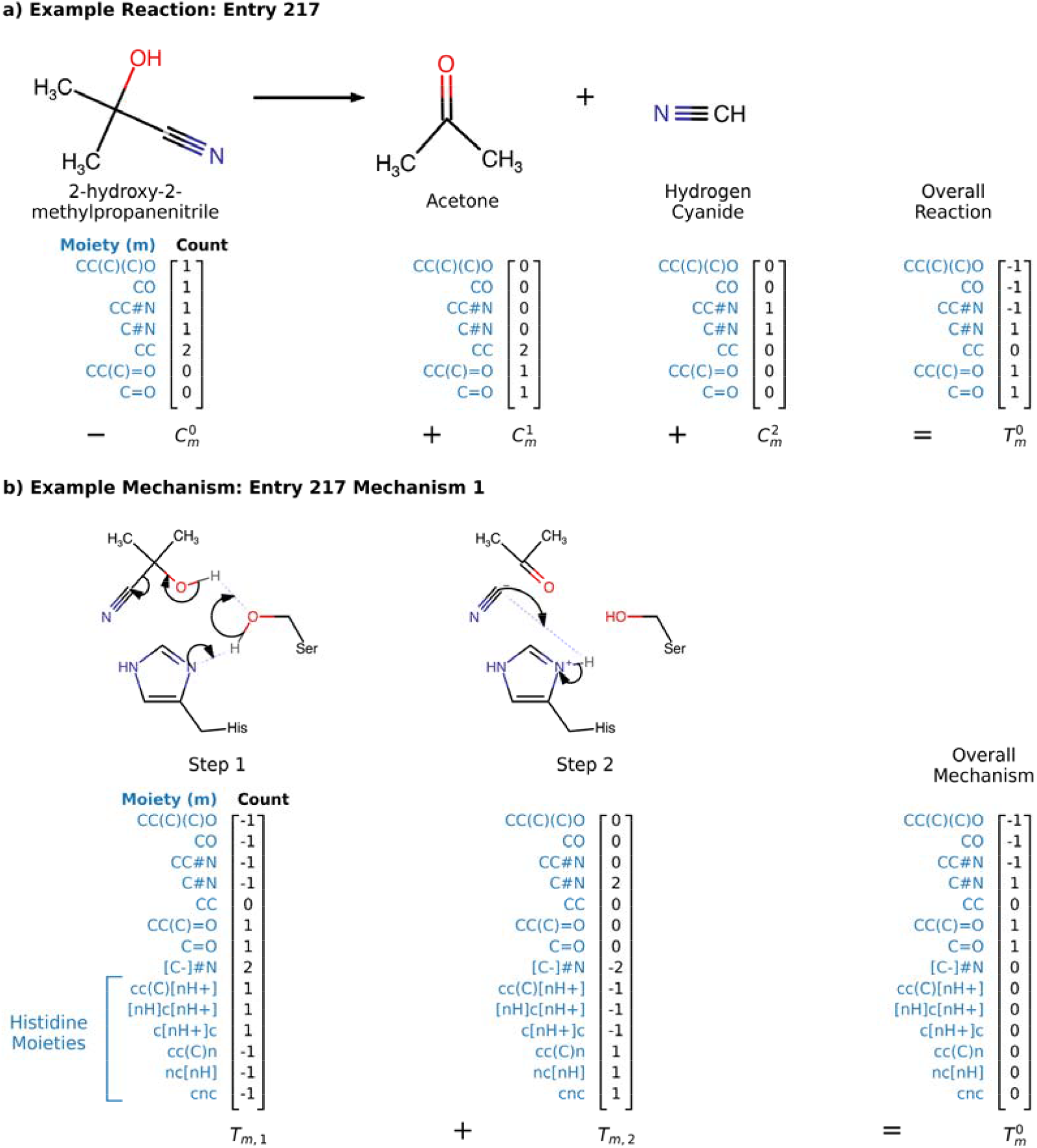
Example entry from M-CSA Database. a) Shows how the reaction is encoded by adding the moiety counts of the products and subtracting the substrates. b) Shows how the complete reaction is decomposed into elementary steps, each encoded using specific moieties. Included in the moiety description are enzyme residues (i.e., Histidine herein) that participate in some of the elementary steps.

### Benchmarking of MechFind on M-CSA entries

Having established a curated set of 4,091 unique elementary rules from the M-CSA database [21], we first performed a comprehensive validation test to assess MechFind’s ability to recapitulate the known mechanisms from the 661 M-CSA entries that could be used as input. This analysis excludes 73 of the 734 reactions because they result in no net change to the moiety fingerprint. For each entry, MechFind was tasked with finding the most parsimonious (i.e., fewest steps) mechanism using only the overall reaction stoichiometry as input.

The framework recovered the correct mechanism (as curated in the M-CSA database) as the top-ranked prediction (rank 1) in 429 of the 661 cases (Figure 2). This means simply by invoking parsimony the correct mechanism is recovered in nearly two thirds of cases (64%). The recovery rate rose to 78% if the correct solution was among the top three predictions and reached 85% of times within the top ten predictions. This implies that by invoking the criterion of parsimony and retaining the top handful of predictions provides strong confidence in recovering the correct mechanism.

**Figure 2.**
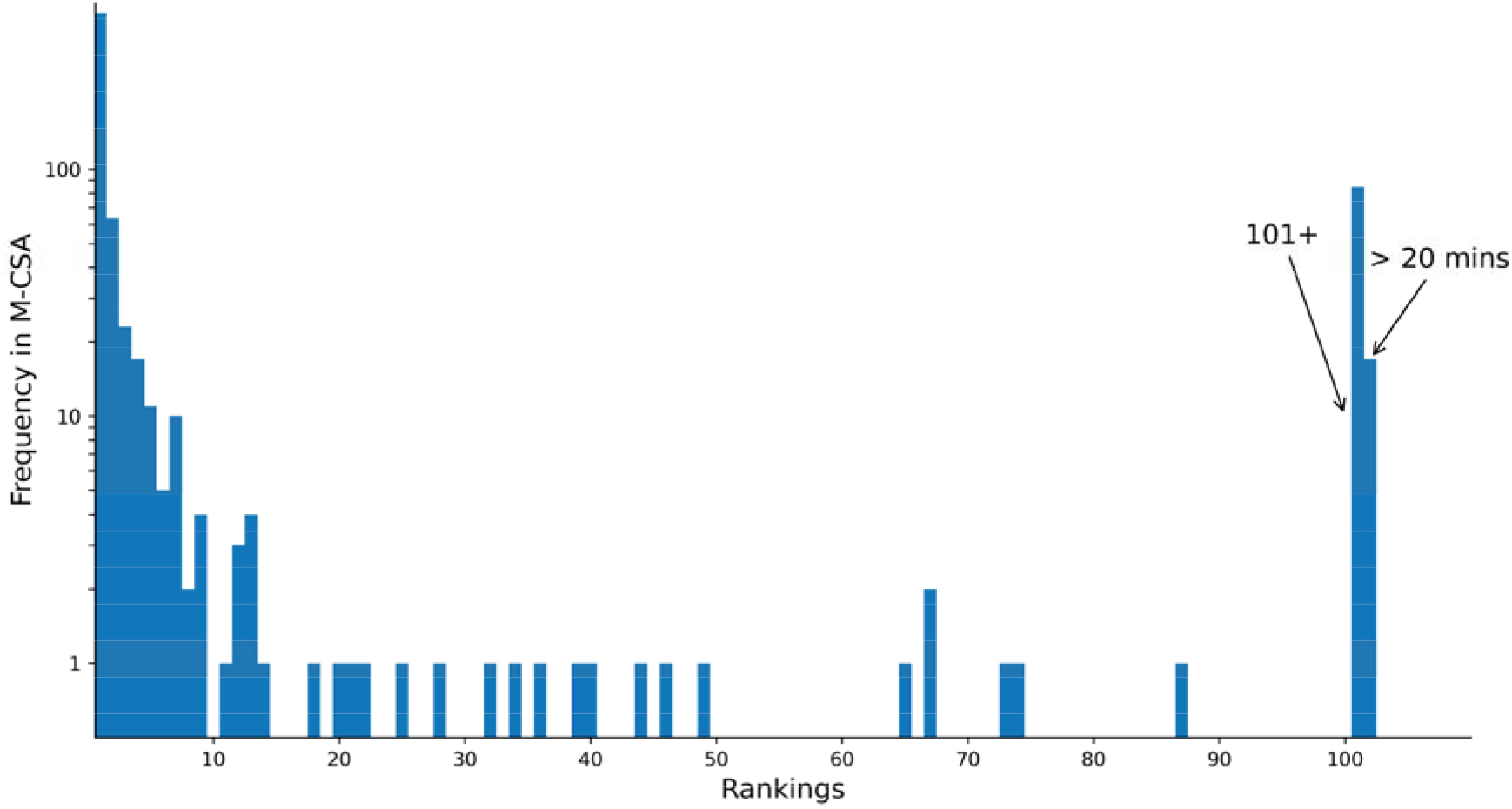
The histogram shows the frequency at which the curated correct mechanism was found at a given rank for the M-CSA entries using the parsimony-based *minRules* and *OrderRules* framework. The “101+” bin represents cases where the correct mechanism was found, but only after more than 100 parsimonious alternatives were generated. The “>20 mins” bin represents cases where the search timed out before a solution was found.

However, in 11% of cases the correct mechanism was not recovered. Inability of recapitulation of the correct mechanism was due to two reasons: either the search for solutions exceeded the 20-minute computational time limit or the framework returned a full list of ten putative mechanisms that did not include the known correct one. This occurs in cases where the principle of parsimony breaks down as the validated mechanism is relatively long even though there exist many shorter, chemically plausible alternatives.

### Independent validation on recently characterized mechanisms unseen in training set

To challenge the framework and demonstrate its potential for genuine discovery, we performed an independent validation using six recently characterized mechanisms absent from the M-CSA training set [31–36]. For each case, we generated the top parsimonious predictions and then re-ranked them using our similarity-scoring method.

The results, detailed in Table 1, show a marked improvement in ranking after applying the similarity based scoring. For example, the correct mechanism for methylisocitrate lyase improved from rank 2 to become the top-ranked prediction. Similarly, the mechanism for glycosaminoglycan lyase saw a significant improvement from rank 8 to rank 4. In four of the six cases, the similarity re-ranker either improved the rank or maintained an already high rank. Crucially, the method never performed worse than the parsimony baseline and successfully identified the correct mechanism within the top 6 predictions for all tested enzymes.

**Table 1.**
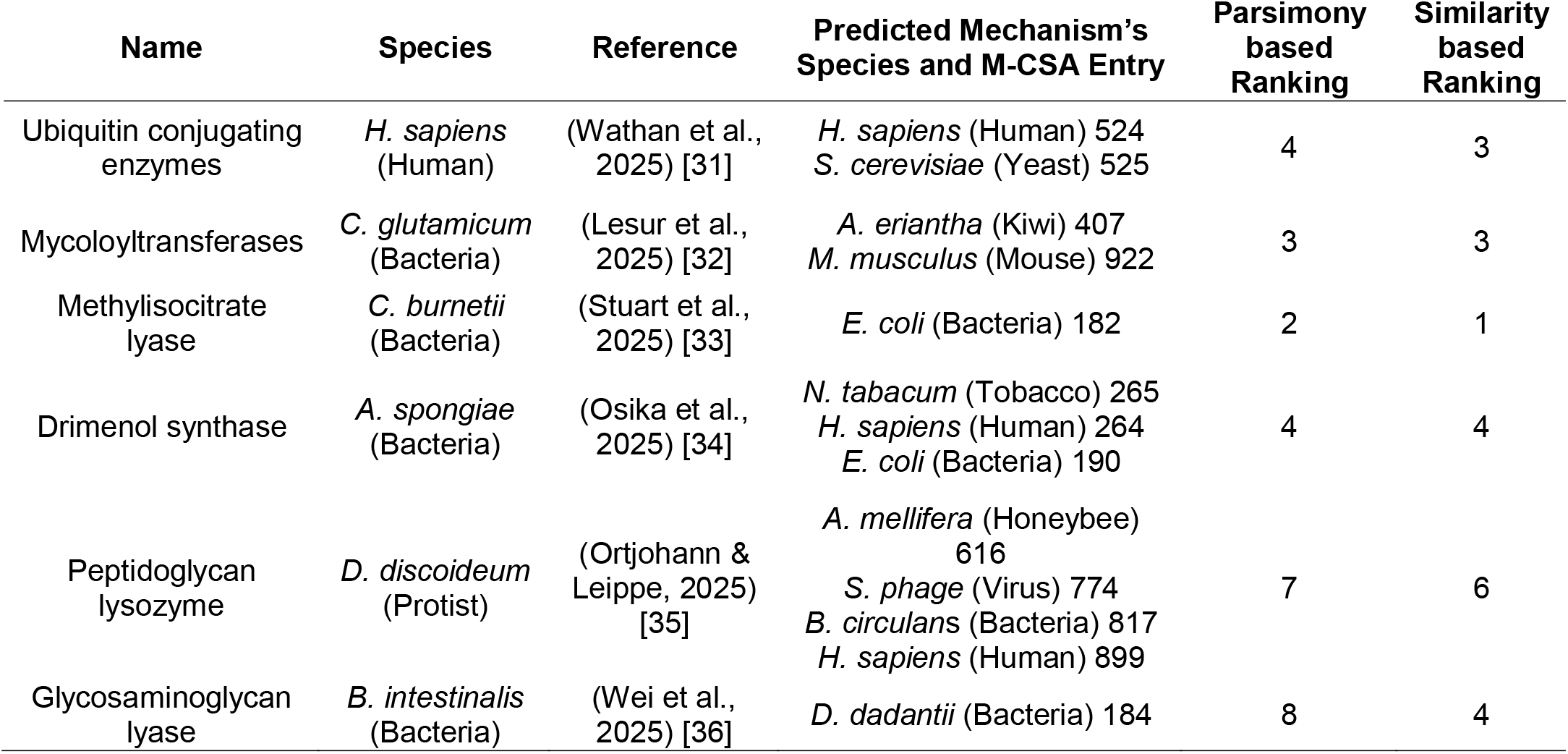
Performance of MechFind on an independent set of mechanisms from novel enzymes.

Notably, the correct mechanistic steps were often constructed by combining elementary rules derived from enzymes in distantly related organisms, highlighting that the fundamental chemical logic of elementary step catalysis can be interoperable across species. For instance, the correct mechanism for a Peptidoglycan lysozyme from the protist D. discoideum [35] was assembled using a combination of elementary rules derived from enzymes in honeybee (*A. mellifera*), a virus (*S. phage*), a bacterium (*B. circulans*), and humans (*H. sapiens*) (see Table 1). Similarly, the mechanism for a human ubiquitin-conjugating enzyme [31] was correctly predicted using rules from both human and yeast acetyltransferases (Figure 3). Although MechFind has no explicit knowledge of ubiquitin or its specific chemistry, it correctly identified the reactive moieties involved—a thioester on acetyl-CoA and a lysine residue’s amine group—and applied a known transformation rule to predict the correct product connectivity. This ability to recognize analogous chemical patterns in vastly different molecular contexts is a key strength of the moiety-based abstraction. A detailed description of the mechanism predictions are shown in Table 1. While most correct mechanisms predictions were among the top 3 predictions in terms of parsimony. The lowest-ranked predicted mechanism was for Glycosaminoglycan lyase (i.e., rank 11). The fact that the correct answer was found within a relatively narrow range of predictions provides confidence that MechFind can serve as a reliable hypothesis-generation tool for novel enzyme mechanism of action.

**Figure 3.**
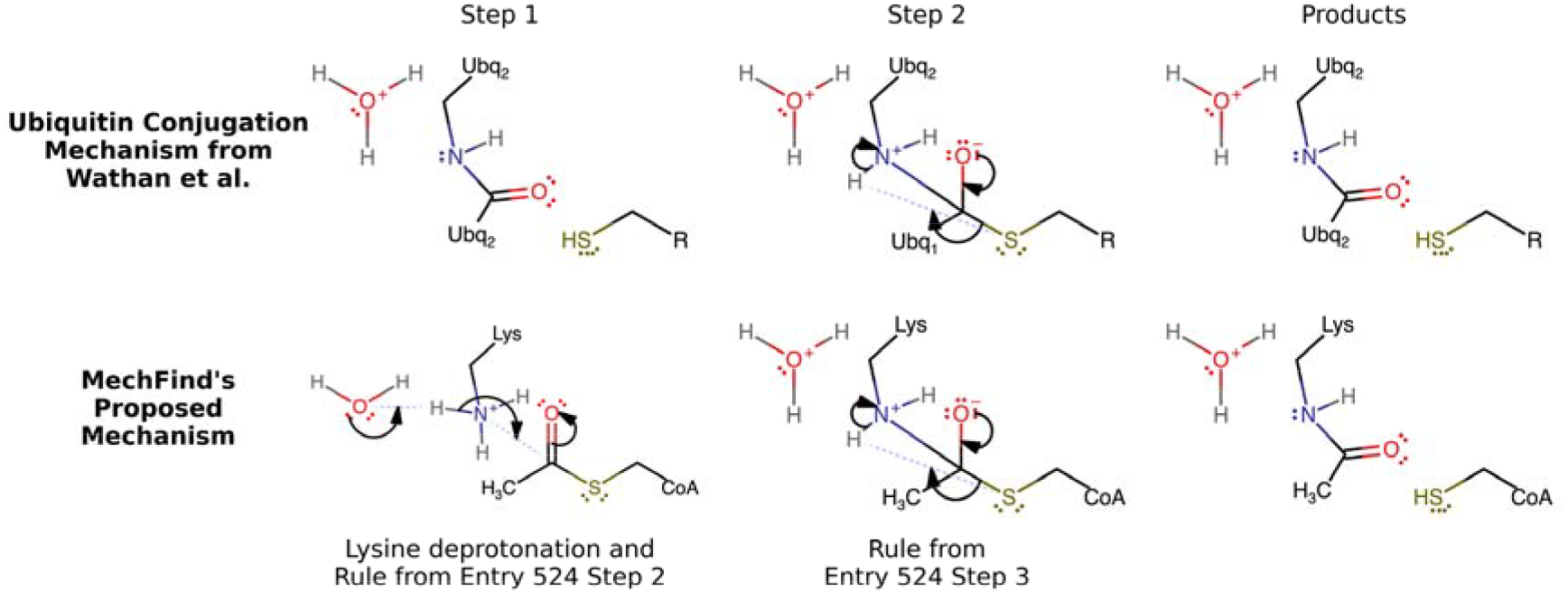
A comparison of the ubiquitin conjugation mechanism from Wathan et al. and a proposed mechanism from MechFind. The proposed mechanism is taken from an acetyltransferase enzyme. Here, we find analogous structures from the ubiquitin mechanism that originate from a lysine residue and a thioester group on acetyl-CoA.

### Large-scale mechanistic annotation of the Rhea and MetaNetX databases

Having established its predictive power and generalizability, we next deployed MechFind to address the growing gap between reaction entries and provided mechanisms in popular reaction databases. We processed 28,412 elementally balanced reactions from the Rhea database [25] and 22,461 from the MetaNetX database [26], the vast majority of which had no prior mechanistic annotation. The outcomes of this high-throughput screen are summarized in Table 2. MechFind successfully generated at least one putative, parsimonious mechanism for 18,100 (63.7%) of the Rhea entries and 14,199 (63.2%) of the MetaNetX entries. This effort represents a more than 20-fold increase in the number of biochemical reactions with available mechanistic hypotheses over prior efforts, creating a public repository that could significantly expand the mechanistic map of known enzyme chemistry. In comparison, the previous state-of-the-art tool, MechSearch, found mechanisms for only 942 (11%) of the reactions it processed from the Rhea database. All predicted mechanisms are available in Supplementary Materials file B.

**Table 2.**
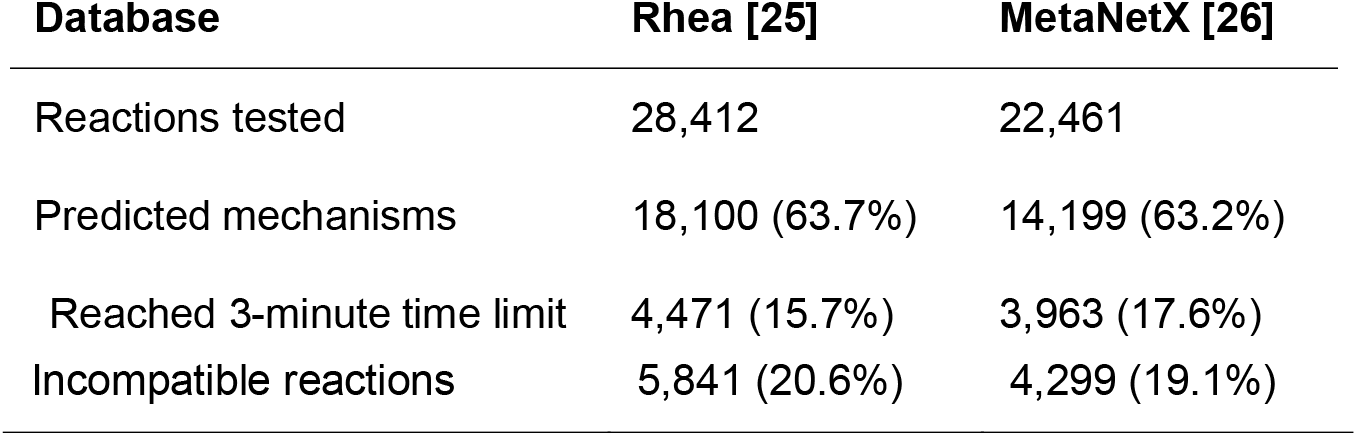
Performance of MechFind on Rhea and MetaNetX Database.

This analysis also helped to identify some of the current limitations of the MechFind framework. As many as 15.7% of Rhea reactions [25] processed by MechFind did not converge within the time limit. While the *minRules* optimization formulation repeatedly proposed sets of elementary steps, the *OrderRules* formulation failed to find a feasible ordering causing the search to exhaust the time limit without a solution. For the 20.6% of reactions where a mechanism could not be constructed, the limitation lies in the scope of our M-CSA-derived rule set [21], as these reactions involve moieties absent in the training data. Figure 4 shows four such examples from the Rhea database, including cyanate (Rhea:11121) and benzyl isothiocyanate (Rhea:10005). While MechFind cannot currently process these reactions, this analysis highlights a clear path for expansion: curating mechanisms involving these out-of-scope moieties and adding them to MechFind’s rule set would directly broaden the applicability of our framework to new classes of enzymatic transformations.

**Figure 4.**
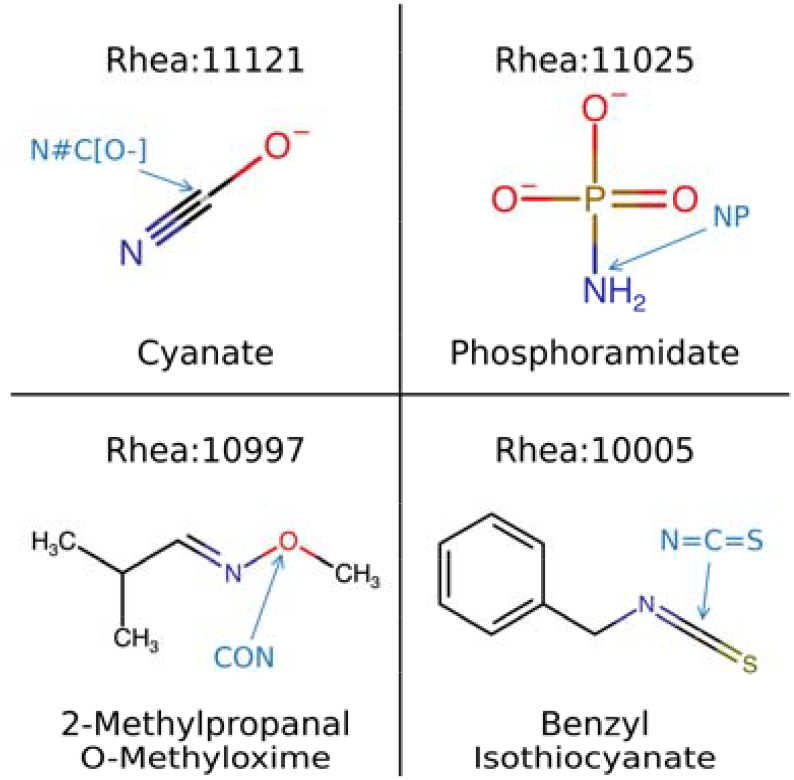
These four compounds are involved in reactions for which MechFind could not propose a mechanism. Each contains a central moiety (highlighted in blue text) that is not represented in the M-CSA-derived rule set, such as cyanate (N#C[O-]), phosphoramidate (NP), 2-methylpropanal O-methyloxime (CON), and Benzyl isothiocyanate (N=C=S).

### Comparisons with existing computational tools

To contextualize the performance of our framework, we conducted a direct comparison with MechSearch [27], the most similar prior method for computational mechanism prediction. This comparison aimed to quantify the differences in input requirements, underlying data, and large-scale performance. The most salient features and outcomes of this analysis are summarized in Table 3. The most fundamental difference lies in the Required Inputs. MechFind requires only reaction stoichiometry, making it applicable to any elementally balanced reaction, whereas MechSearch’s need for user-supplied active site residues limits its use to enzymes with known or hypothesized structures. MechFind leverages a more comprehensive set of rules derived from 734 curated M-CSA mechanisms [21], nearly double the 368 used by MechSearch. This translates to a disproportionately higher recovery performance by MechFind. On a comparable set of reactions from the Rhea database, MechFind processed over 28,000 reactions with a Success Rate of 64%. This represents a nearly six-fold improvement over the 11% success rate reported for MechSearch on a smaller subset of the database.

**Table 3.** Feature and Performance Comparison of MechFind with MechSearch.

| **Feature** | **MechFind (This work)** | **MechSearch [27]** |
| --- | --- | --- |
| **Required Inputs** | Reaction Stoichiometry Only | Stoichiometry + User-Supplied Active Site Residues |
| **External Data Requirement** | None | Protein structure or active site hypothesis |
| **Rule Set Source** | 734 curated M-CSA mechanisms | 368 M-CSA mechanisms |
| **Large-Scale Application** | Rhea Database | Rhea Database |
| **Reactions Processed** | 28,412 | 8,805 |
| **Success Rate** | 64% (18,100 / 28,412) | 11% (942 / 8,805) |
| **Output Format** | Ranked list of top-10 parsimonious mechanisms | Set of putative mechanisms |

### Application of MechFind to explore mechanistic diversity

Beyond identifying a single most-parsimonious mechanism, MechFind can be used as an exploratory engine to map the network of plausible catalytic pathways for a given transformation. We applied this capability to a canonical esterase reaction, constructing the mechanistic network shown in Figure 5 by compiling the unique elementary steps from the top-ten predicted mechanisms. This network reveals a landscape of competing catalytic strategies, such as the formation of a serine covalent intermediate versus a direct nucleophilic attack by water, demonstrating that multiple distinct mechanistic routes are often chemically plausible for a single overall reaction.

**Figure 5.**
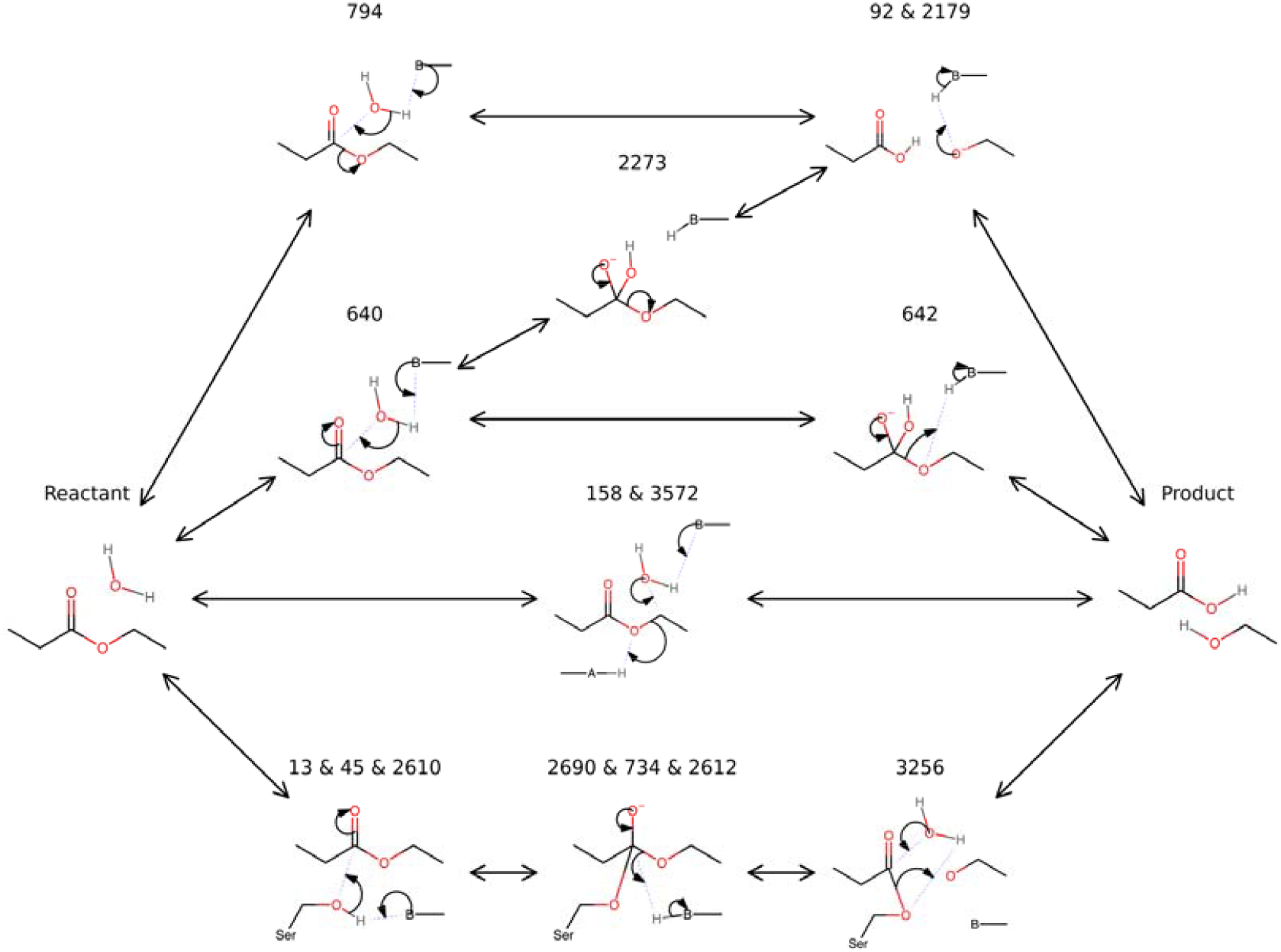
The mechanistic network for a canonical esterase reaction was constructed by compiling the top-ten most parsimonious mechanisms predicted by MechFind. Each unique elementary rule shown has its own number; for the steps that have multiple numbers, multiple different general bases have been observed, these were combined for simplicity. General bases are denoted with the letter B, whereas general acids are denoted with the letter A. These general items could be water, oxonium, hydroxide or any of the electrically charged amino acids.

This landscape of mechanistic diversity presents a powerful opportunity for de novo enzyme design. The bridge between a predicted mechanism and a functional enzyme lies in the principle of transition state stabilization [19]. Each elementary step in the network proceeds through a unique transition state. By using computational chemistry methods to find the lowest-energy path between the predicted intermediates of a given step, the highest-energy point can be identified as the putative 3D transition state structure. These structures are the precise, actionable targets required by advanced generative models like RFdiffusion All-Atom to build a stabilizing protein scaffold (Krishna et al., 2024). MechFind, therefore, provides a menu of these potential transition states, transforming the abstract challenge of “designing an esterase” into a series of concrete, testable engineering tasks, such as, “design a protein active site that stabilizes the tetrahedral transition state formed by Rule 640.” In this way, MechFind’s ability to map the mechanistic landscape provides a rich foundation of parallel hypotheses to guide the rational design and engineering of novel enzymes with tailored catalytic functions.

## Discussion

In this work, we introduced MechFind, a computational framework designed to address the significant and growing gap in mechanistic information for catalogued biochemical reactions. By leveraging a curated set of elementary chemical rules from the M-CSA database [21], MechFind predicts detailed, multi-step enzyme mechanisms using only overall reaction stoichiometry as input, employing a hybrid approach that combines mixed-integer linear programming with a similarity-based re-ranker that scores candidate mechanisms against our entire database of validated mechanisms. The framework’s efficacy was first established by recapitulating the correct mechanism within the top ten predictions for 85% of the 661 known M-CSA entries, with the correct mechanism identified as the top-ranked, most parsimonious solution in 64% of cases. Its capacity for genuine discovery was then demonstrated on an independent validation set of six novel enzymes, where it successfully identified the correct mechanism in all cases, often by combining chemical rules from distantly related organisms. When deployed at scale, MechFind generated over 18,000 putative mechanisms for 64% of the 28,412 reactions tested from the Rhea database [25], representing a more than 20-fold increase in the number of reactions with available mechanistic hypotheses and a nearly six-fold performance improvement over existing tools. Finally, we demonstrated MechFind’s utility not just as a prediction tool, but as an exploratory engine for generating a comprehensive network of testable hypotheses about the diverse catalytic strategies available for a given reaction, as shown for a canonical esterase.

The primary advance of MechFind lies in its ability to bypass the main bottleneck that has limited previous computational approaches. Methods such as MechSearch [27] and EzMechanism [28] require user-supplied active site residues or high-resolution protein structures, respectively, restricting their application to a small subset of well-characterized enzymes. This prerequisite makes the high-throughput analysis of entire databases infeasible. By removing this requirement, MechFind enables, for the first time, the high-throughput mechanistic annotation of entire reaction databases. This capability begins to bridge the “mechanism gap” between the tens of thousands of known biochemical reactions and the fewer than one thousand with detailed catalytic steps. The resulting library of over 18,000 high-confidence mechanistic hypotheses provides a critical starting point for downstream rational enzyme engineering efforts. It fundamentally changes the paradigm from a low-throughput, hypothesis-driven process, where a researcher must first pinpoint which residues are involved, to a high-throughput, data-driven discovery process that formalizes the generation of hypotheses.

Our large-scale application of MechFind yielded non-obvious insights into both the conservation of enzymatic strategies and the nature of the challenges that remain. The independent validation results strongly suggest the existence of a universal “chemical grammar” that is conserved across species. For example, the mechanism for a Mycoloyltransferase from the bacterium C. glutamicum was correctly constructed using elementary rules derived from enzymes in a kiwi plant and a mouse (Table 1). Similarly, the mechanism for a human ubiquitin-conjugating enzyme was correctly predicted using rules from yeast acetyltransferases (Figure 3). These findings demonstrate that fundamental catalytic principles are modular and can be recognized and redeployed by the model, regardless of the evolutionary context or the overall molecular scaffold. This insight validates our core moiety-based abstraction, proving that the local chemical environment is a powerful descriptor for predicting enzymatic reaction mechanisms across the kingdoms of life. It implies that the vast diversity of enzymatic function arises not from an unbounded set of chemical possibilities, but from the recombination of a relatively small set of elementary reaction rules.

While this work demonstrates a significant step forward, it is essential to acknowledge the framework’s current limitations, which define clear directions for future development. Our analysis of the 36% of Rhea [25] reactions for which no mechanism was found reveals that the problem is twofold. We identified that for 16% of reactions, the challenge stems from a limitation in our decomposed optimization strategy, where the *minRules* formulation proposes rule sets for which *OrderRules* cannot find a valid sequence within the time limit. This points to the need for more efficient optimization algorithms or heuristics for particularly large and complex substrates. In addition, we found that for 21% of reactions, the limitation is the scope of our M-CSA-derived rule set [21], which lacks the necessary chemical moieties to describe the transformation. This highlights that the model’s predictive power is fundamentally bound by its library of known chemistry; it can combine known steps in novel ways but cannot invent entirely new transformations. The clear path forward is to continue expanding the rule set by curating more diverse mechanisms. Furthermore, our current implementation deliberately aggregates stereoisomers. Given training data, the incorporation of stereo-specific rules would unlock more complex mechanisms for MechFind.

Perhaps the most significant impact of this work, however, lies in proposing a path to close the loop for automated enzyme design. As established in the introduction, the goal of de novo enzyme design is to create a protein active site that stabilizes the transition state of a desired reaction [19]. The newest generation of protein design tools, such as RFdiffusion All-Atom [11] and ProteinMPNN [12], are very effective at generating protein structures but require a precise structural target to aim towards. The mechanistic hypotheses generated by MechFind provide the putative transition states that these advanced design tools require. This transforms the abstract challenge of “designing an enzyme for this reaction” into the concrete and actionable task of “designing a protein scaffold to stabilize this specific proposed mechanism.” This is particularly powerful when considering the diversity of possible mechanisms (see Figure 5) that MechFind can uncover offering a roadmap of candidate transition states that can serve as targets for parallel design efforts. In conclusion, MechFind provides both a foundational tool and a public resource that significantly expands the map of known enzyme chemistry, paving the way for the next generation of rational enzyme engineering.

## Materials and Methods

### Database Curation and Rule Generation

A prerequisite for building a predictive framework based on chemical transformations is a dataset that is internally consistent and adheres to mass and charge conservation laws. Curation began with the 734 fully annotated mechanisms in the M-CSA database [21], which form the basis of our elementary rule set. Initial analysis revealed that a significant fraction of these entries contained inconsistencies, such as incorrect protonation states, inconsistent stereochemical assignments, or mis-drawn homolytic versus heterolytic bond events. These issues resulted in violations of elemental and charge balance for either individual steps or the overall reaction. To rectify this, we manually curated the entire set of 734 mechanisms. Due to stereochemical ambiguities in most catalogued mechanisms we opted to bypass molecular chirality and simply represent all compounds by their constitutional isomers. Beyond this simplification, other corrections were designed to be as minimal as possible, primarily addressing inconsistencies in protonation states and ensuring that arrow-pushing schemes correctly represented bond reorganizations. In total, 463 entries (63%) required at least one correction. For example, in M-CSA entry 66 (see Figure 6a), the protonation state of the product was inconsistent with the elementary steps, requiring a correction to the overall reaction stoichiometry (Figure 6b). Following the curation process a final, self-consistent set of 3,235 reversible mechanistic steps was assembled. Each step was treated as reversible, creating a forward and backward version, and combined with protonation steps of common moieties to generate the 4,091 unique elementary rules that underpin MechFind. Note that moieties are simply classifications of all non-hydrogen atoms based on their first bonding shell present in the metabolites forming the database. Reaction rules abstract the overall reaction as the gain or loss of specific moieties.

**Figure 6.**
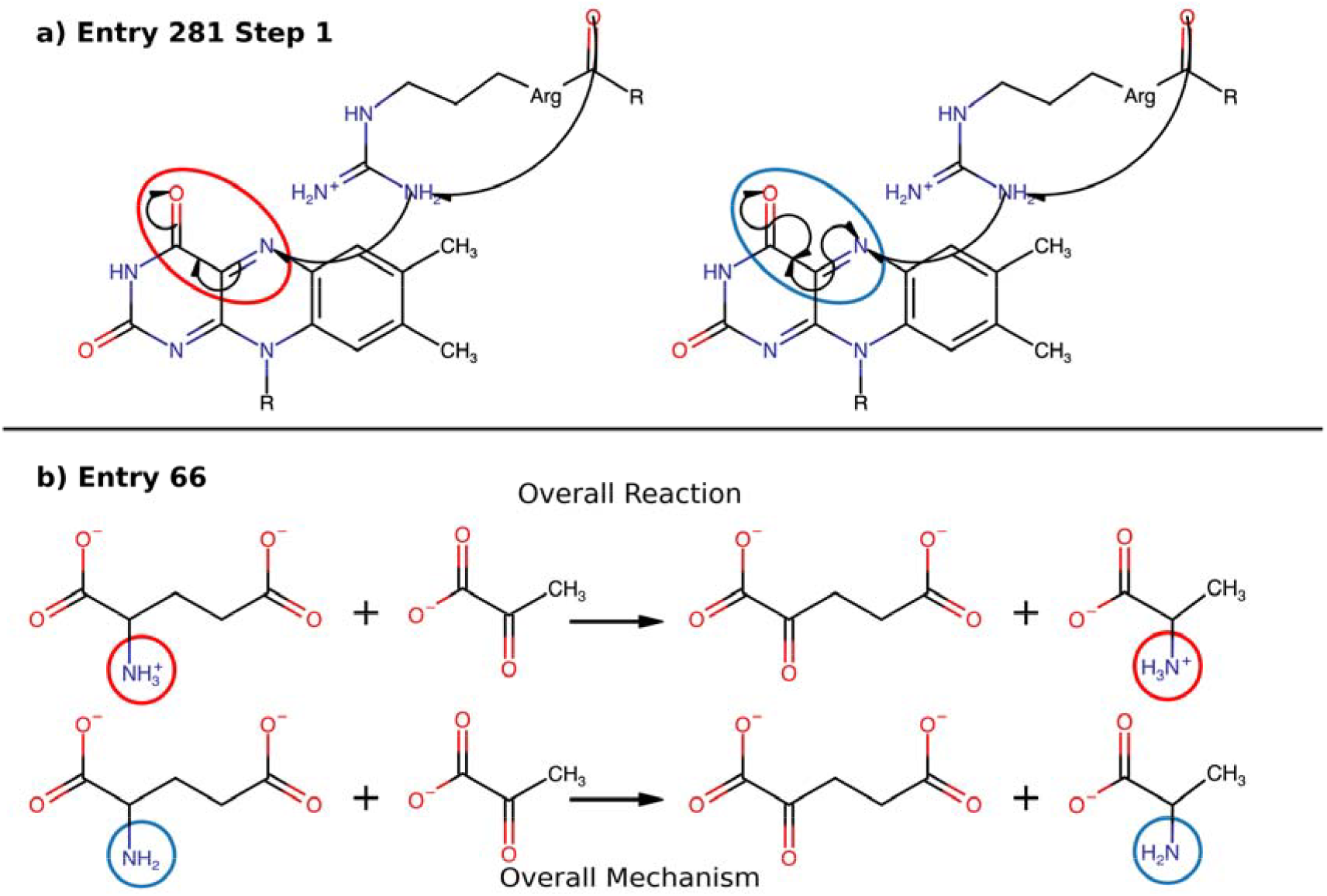
Examples of common adjustments made in the molecules in the M-CSA database with the original state circled in red and their correct state in blue. The overall mechanism is determined by summing all molecules at every step in the mechanism, thereby subtracting the reactants in each step from the products. a) A correction to the mechanism due to homolytic bond formation and cleavage not being correctly depicted using arrow pushing. b) A correction to the overall reaction is required due to the protonation states of the product and reactant being inconsistent with those shown in the overall mechanism in the database.

### Optimization Formulation for Mechanism Prediction

To assemble plausible, stepwise enzymatic mechanisms, the fundamental challenge is to simultaneously identify the necessary set of elementary reaction rules and establish a feasible order. Conceptually, this can be addressed by a single mixed-integer linear programming (MILP) formulation, which we term *minOrderRules*. This formulation minimizes the total number of steps subject to constraints ensuring moiety balance and a valid sequencing order that prevents the use of moieties not present in the reactants or synthesized beforehand.

### minOrderRules

The *minOrderRules* formulation uses integer variable, *y*_*r*_, to represent the number of times a rule *r* is used, and binary variable *z*_*k,r*_ to assign rule *r* to a specific step *k* in the mechanism.

Objective function

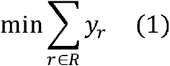

### Subject to

Moiety balance:

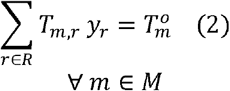

Cumulative sum:

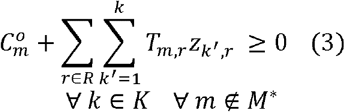

- This objective minimizes the total number of elementary steps, enforcing the principle of parsimony.
- This constraint ensures that the sum of moiety changes from all applied rules equals the net change associated with the overall reaction.
- This key constraint ensures chemical plausibility by preventing the consumption of any substrate, or intermediate moiety before it has been created or was present in the initial reactants. It accomplishes this by ensuring that the cumulative balance of these specific moieties remains non-negative at every step *k* of the mechanism. Here, 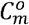 is the initial count of moiety *m* in the substrates.

Crucially, this constraint is applied to all moieties m that are not part of the set *M*^*^, which represents all labeled moieties originating from catalytic residues and cofactors. This distinction is essential because catalytic groups are considered part of the enzyme’s reusable machinery. For example, this allows a protonated histidine to be consumed in one step and a deprotonated form to be generated in another without violating the constraint, as both histidine moieties are in *M*^*^ and thus exempt from this rule. This gives the model the necessary flexibility to utilize the different states of catalytic residues and cofactors as required throughout the mechanism, reflecting their dynamic role in the active site. A complete list of the labeled moieties in *M*^*^ can be found in Supplementary Materials file A.

One rule per step:

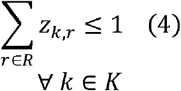

Step utilization limit:

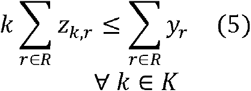

Contiguous steps:

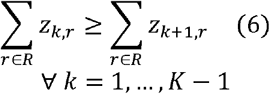

Linking variables:

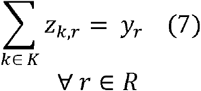

Integer cuts:

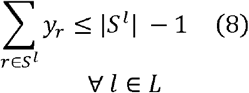

- This enforces that each step *k* in the mechanism makes use of at most one elementary rule.
- This constraint ensures that steps are used only if they are within the total number of steps required for the mechanism. It effectively forces *z*_*k,r*_ to be zero for any step *k* that is greater than the total number of steps ∑*y*_*r*_.
- This ensures that the sequence of steps is contiguous, with no empty (i.e., rule unassigned) steps between steps in use.
- This constraint links the step assignment variable *z* _*k,r*_ to the rule count variable *y*_*r*_, ensuring that the total number of times a rule is assigned to a step equals its total count.
- To generate a ranked list of the top-ten most parsimonious mechanisms, specialized integer cut constraints [38] are used iteratively. After finding a solution *l*, we define *s*^*l*^ as the set of rules *r* with *y*_*r*_ > 0. The corresponding integer cut constraint excludes the prior solution from the set of feasible choices without excluding any other combination. By successively appending integer cut constraints and resolving the optimization problem a ranked list of optima (i.e., first, second, third, etc.) is obtained.

### Decomposed Formulation for Computational Tractability

However, early trials revealed that solving *minOrderRules* formulation is computationally taxing for all but the simplest reactions, with solution times often taking several minutes per candidate mechanism. To create a high-throughput and scalable tool, we decomposed this complex task into two separate MILP problems: *minRules*, which first identifies the needed rules, and *OrderRules*, which subsequently finds a feasible ordering. In the *OrderRules* formulation, the total number of steps in the mechanism is no longer a variable to be optimized but is instead a fixed integer, defined by the sum of the *y*_*r*_ values in the solution set *s*^*l*^ provided by *minRules*. This dramatically reduces the combinatorial search space for the second, more complex ordering problem. Consequently, the constraints for ensuring step contiguity and limiting step utilization (Equations 5 and 6) are no longer required. Conversely, in *minRules* formulation the maximum possible number of rules must be directly specified (Equation 11).

### minRules

This first formulation, *minRules*, identifies the minimal set of rules required to satisfy the overall reaction stoichiometry.

Objective function:

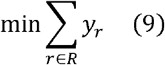

Subject to:

Moiety balance:

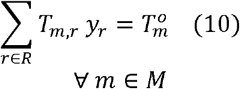

Maximum number of rules:

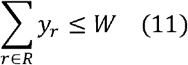

Integer cuts:

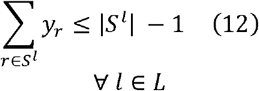

- This objective function is identical to Equation (1).
- This constraint is identical to Equation (2).
- This limits the total number of elementary steps to a user-defined value *w* (herein set to 20).
- This constraint is identical to Equation (8) and is used to generate multiple unique sets of rules.

### OrderRules

For each set of rules *s*_*l*_ found by *minRules*, this second formulation determines if a feasible sequence exists. Therefore, its objective function is a dummy function that plays no role in the solution.

Subject to:

Cumulative sum:

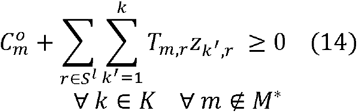

One rule per step:

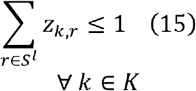

Each rule must be used:

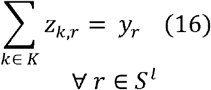

- This constraint is analogous to Equation (3).
- This constraint is analogous to Equation (4).
- This constraint, analogous to Equation (7), ensures every rule from the *minRules* solution is used in the sequence.

### Similarity-Based Re-ranking

To improve the recovery rate of correct mechanisms beyond simple parsimony, we first attempted to train a deep neural network to re-rank the top candidates. This approach, however, could not improve upon the initial parsimony-based rankings (see Supplemental File A for details). We therefore developed an alternative method to re-rank the top-ten candidate mechanisms generated by the *minRules* and *OrderRules* formulations. This successful method scores each candidate based on its similarity to the known mechanisms in the M-CSA database [21].

The re-ranking process begins with the list of the top-ten most parsimonious mechanisms for a given reaction. Each of these candidate mechanisms undergoes a pairwise comparison against every validated mechanism in the curated M-CSA database. The similarity is quantified using the “unordered” score variant described by Ribeiro et al. [39], which is calculated over the sets of elementary chemical steps that constitute each mechanism and ranges from 0 (no shared steps) to 1 (identical sets of steps). Each candidate mechanism is then assigned a single score corresponding to the maximum similarity value obtained from all its pairwise comparisons, which represents its similarity to its closest known analog in the database. Finally, the ten candidates are re-ranked in descending order of their assigned similarity scores, with the candidate having the highest score ranked first.

### Data Acquisition from Public Databases

To apply MechFind on a large scale, we integrated reaction data from Rhea [25] and MetaNetX [26]. Rhea supplies a standardized list of 34,576 biochemical reactions. Because our current framework does not differentiate between stereoisomers, 4,016 reactions involving epimerase and racemase enzymes were removed. An additional 2,148 reactions that were not stoichiometrically balanced were also removed, reducing the total to 28,412 reactions from Rhea for application in MechFind. A similar process was performed for the MetaNetX database, which contains 23,586 balanced non-transport reactions. After removing 1,125 reactions involving epimerases and racemases, a final set of 22,461 reactions from MetaNetX was used for testing. A significant overlap exists between these two curated datasets, with 9,318 (41%) of the reactions in our final MetaNetX set also being present in the Rhea set. The distribution of these reactions across the seven main Enzyme Commission (EC) classes is shown in Figure 7, highlighting that both databases are predominantly composed of oxidoreductases (EC 1), transferases (EC 2), and hydrolases (EC 3). Notably, a significant fraction (i.e. 57% in Rhea and 28%, in MetaNetX) of both databases do not have any EC number associated with it. A complete list of the reactions included from both databases is available in Supplemental Files B.

**Figure 7.**
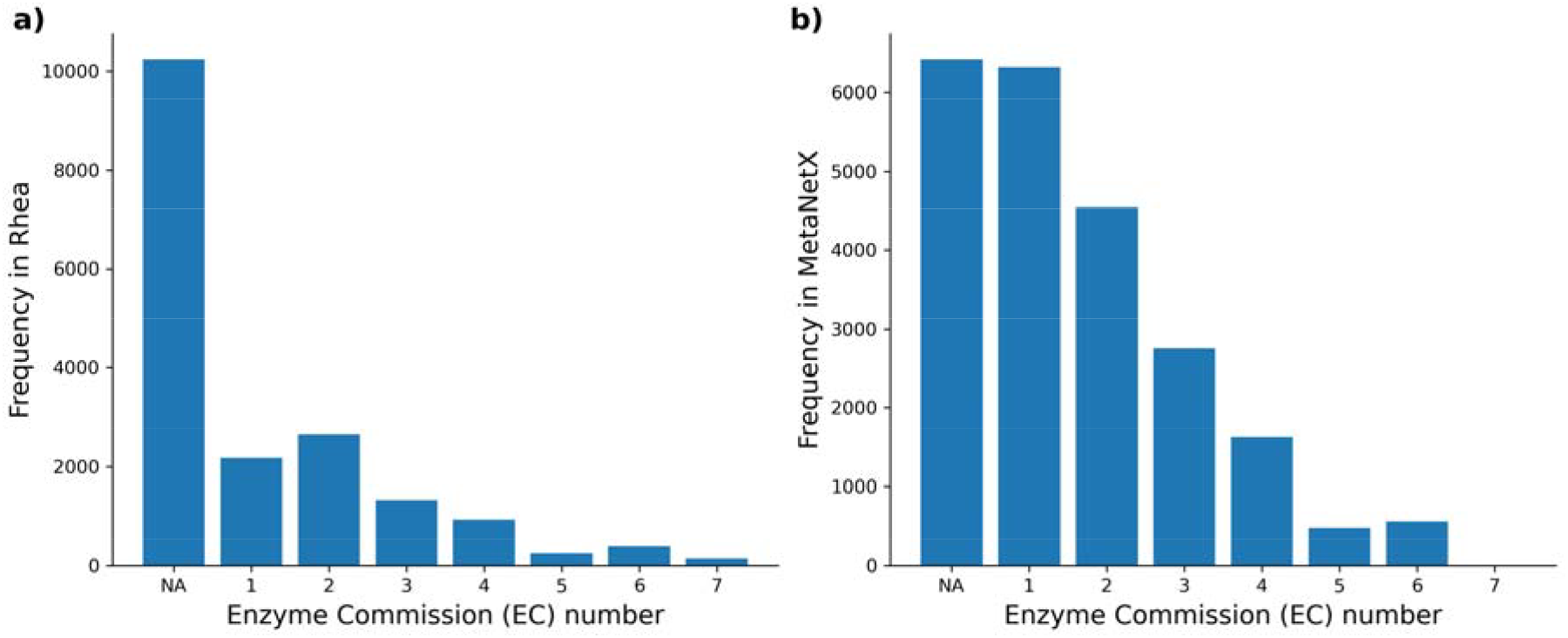
The bar charts show the frequency of reactions across the seven main EC classes for (a) the 28,412 reactions used from the Rhea database and (b) the 22,461 reactions used from the MetaNetX database. The ‘NA’ category includes reactions without an assigned EC number. Both databases are dominated by oxidoreductases (EC 1), transferases (EC 2), and hydrolases (EC 3), reflecting the prevalence of these catalytic functions in known metabolism.

## Supporting information

Supplemental File A, more detail on the curation process and attempts at re-ranking

Supplemental File B, solutions for reactions in Rhea and MetaNetX database

## Data Availability

The curated M-CSA mechanisms, elementary rules matrix (Unique_Rules.csv), and arrow environment data (M-CSA_arrow_rules_r0.json) used in this study are available in the GitHub repository. The full list of reactions from the Rhea and MetaNetX databases that were processed are available in the Supplementary Information. All other data are available from the corresponding author upon reasonable request.

## Code Availability

The source code for the MechFind framework and the Jupyter notebook to reproduce the example analysis are publicly available on GitHub at https://github.com/maranasgroup/MechFind

## Acknowledgments

This material is based upon work supported by the Center for Bioenergy Innovation (CBI), U.S. Department of Energy, Office of Science, Biological and Environmental Research Program under Award Number ERKP886. Any opinions, findings, and conclusions or recommendations expressed in this publication are those of the author(s) and do not necessarily reflect the views of the U.S. Department of Energy. This work was also supported by the U.S. National Science Foundation funded Molecule Maker Lab Institute (MMLI), award number 2019897 supported by National AI Research Institutes Program of the Directorate for Computer and Information Science and Engineering (CISE), in collaboration with the Division of Chemistry (CHE) and the Division of Chemical, Bioengineering, and Environmental Transport Systems (CBET) awarded to CDM. This publication was supported by the National Institutes of Health (NIH) Training Grant Number 5T32GM149417. The funders had no role in study design, data collection and analysis, decision to publish, or preparation of the manuscript. The authors of this work recognize the Penn State Institute for Computational and Data Sciences (RRID:SCR_025154) for providing access to computational research infrastructure within the Roar Core Facility (RRID: SCR_026424).

## Author Contributions

Austin D. Hartley – Conceptualization, Methodology, Software, Validation, Formal analysis, Investigation, Resources, Data Curation, Writing (Original Draft), Visualization

Vikas Upadhyay – Conceptualization, Software, Supervision Veda Sheersh Boorla – Conceptualization, Software, Supervision

Costas D Maranas – Conceptualization, Writing (Review & Editing), Supervision, Project administration, Funding acquisition

