## Supplemental File A, more detail on the curation process and attempts at re-ranking for "MechFind: A computational framework for de novo prediction of enzyme mechanisms"

Supplementary Materials

Contents

1. Curation and Encoding of M-CSA Mechanisms

1. Encoding Arrow-Pushing Schemes

2. Curation Process

3 Labeling codes for catalytic groups

2. Neural Network Re-ranking Analysis

1. Methods - Neural Network Architecture

2. Results and Discussion

S1. Curation and Encoding of M-CSA Mechanisms

S1.1. Encoding Arrow-Pushing Schemes

To convert the graphical arrow-pushing diagrams from the M-CSA into a machine-readable format for our optimization framework, we developed a set of rules to systematically interpret the change in charge and bond order for the atoms involved in each arrow. These rules, summarized in Table S1, define how double-headed (two-electron) and half-headed (one-electron) arrows are translated into changes in the local chemical environment. This encoding allows each elementary step to be represented as a vector of moiety changes.

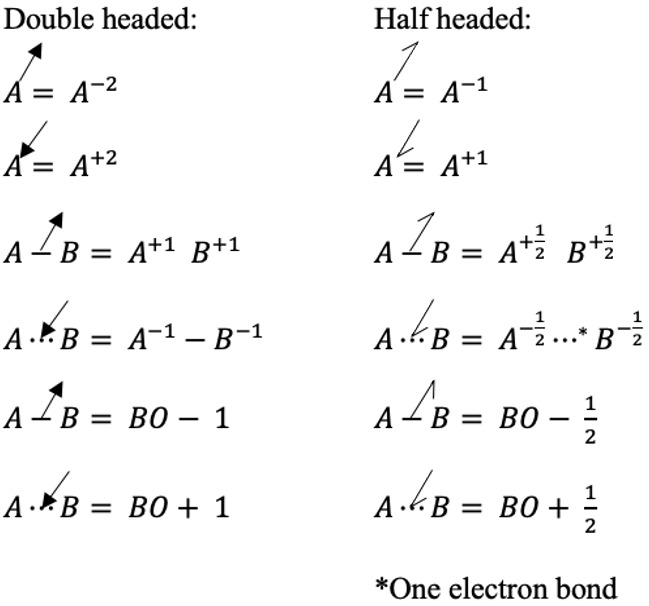

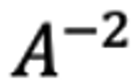

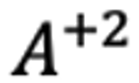

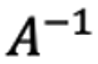

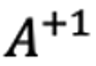

Figure S1. Rules for translating arrow-pushing notation into changes in charge and bond order. The table details the effect of double-headed (2e⁻) and half-headed (1e⁻) arrows on atoms (A, B) and bond orders (BO). A half-headed arrow pointing to a bond indicates the formation of a one-electron bond.

S1.2. Curation Process

As described in the main text, the 734 mechanisms from the M-CSA database underwent a manual curation process to ensure internal consistency. This involved correcting inconsistencies in protonation states, stereochemical assignments, and mis-drawn bond events to ensure that both individual steps and the overall reaction were elementally and charge-balanced. All Changes made to each step are documented below.

Entry 5

extra R group on the mechs reactant

5 1 2

5 1 3

5 1 4

5 1 5

Entry 20

first step creates the wrong FADH state, 2 e- need to flow from a donator, in this case its known to be menaquinol heme only transfers single e- so extra steps are needed the first step is replaced with a menaquinol deprotonation the second step transfers the e- and N5 is radicalized the third step deprotonates the menaquinol again and transfers an e- allowing for N5 to be protonated as it is in the following step steps 2,3, and 4 are shifted up by two to fit the new steps

20 1 1

20 1 2

20 1 3

20 1 4

20 1 5

20 1 6

Entry 37

arrow from oxygen should go from Fe and the Fe N bond to the Fe O bond changing the Fe charge back to +3 and creating the double bond the product has an extra oxygen on it and the heme is not reset this is the best I could fix the reaction is also unbalanced

37 1 2

37 1 3

Entry 62

the bond from the cobalt to its base tail is a binding bond in the first step but a covalent bond in the last step, and the charge on the cobalt is -2 for every step but 1, step one is updated to have the covalent bond and changes the cobalt’s charge to -2

62 1 1

Entry 68

the FAD reactant is not radicalized or protonated properly I assumed OH3+ protonated both the N1 and N5 positions on the FAD

68 1 2

68 1 3

Entry 90

step 5 deletes the C5 carbon on adenosine

90 1 5

Entry 102

heme iron in step 3 should be -1 charge, neutral in step 4, and -1 charge in step 5, also the FAD is missing its radical in step 4 so it is added back

102 1 3

102 1 4

102 1 5

Entry 105

step 2 Mo should be charge +5 also Mo is missing =O atom, step 2 Mo should gain another e- from the new bond with Glu that is later destroyed, step 3 Mo losses an e- to the 1,4-benzoquinones this is a overall reactions replacement for the e- acceptor, step 4 Mo losses the last e- resetting, and hydroquinone is formed the proton donors were assumed to be OH3+

105 1 2

105 1 3

105 1 4

Entry 107

binding bonds cause most of the conflict here they do not behave consistently across the mechanism

if you assume all of the binding bonds do not behave like covalent bonds the Cu does not seem to have to change charge in the mechanism the Mo charge is correct though out if you assume that OH- did not form a covalent bond in step 4, step 5 and 6 do not properly radicalize or protonate the quinones reactant assumed OH3+ protonated

107 1 2

107 1 3

107 1 4

107 1 5

107 1 6

Entry 113

the reactant should start protonated on the nitrogen

113 1 1

Entry 114

FAD reactant is not radicalized or protonated correctly assumed OH3+ was the proton donor in step 4 Marvin does not allow for a radical half arrow to go from an atom to a bond that is nonadjacent to that atom to fix this the half arrow was snaked from one of the nitrogen atoms towards the new double bond meeting with the initial radical on the FAD, step 6 makes a 6 carbon bond

114 1 3

114 1 4

114 1 6

Entry 121

The charge on molybdenum after step 1 should be +5 due to the Mo O double bond breaking to a single and the double headed arrow moving to the Mo the bond breaking adds +1 charge to both atoms and the double headed arrow means that 2 e- add their -1 charge to the Mo adding to net -1 charge on the Mo steps 2,3,and 4 were updated to be consistent with this curious if a Mo+4 atom is ever experimentally observed if so then this mechanism is wrong

121 1 2

121 1 3

121 1 4

Entry 123

to better fit with other rules the general donor is replaced with a 4 atom iron sulfur clusters

123 1 1

123 1 2

Entry 124

step 1 bonds the iron to the O=O, step 2 creates the second bond from Fe to O=O while grabbing an e- from the copper so it better aligns with later steps, steps 3-8 are changed to only maintain the -2 charge on the iron and +2 charge on the copper except in step 8 when the copper is reduced from step 7, step 9 should take e- from the Fe O bond not the atom thus oxidizing the iron allowing it to be reset in step 10, step 12 should use the last e- to reset the copper back to a +1 state the protonation chains in steps 3,4,6,7,9,10, and 12 were simplified to reduce substructure matches

124 1 1

124 1 2

124 1 3

124 1 4

124 1 5

124 1 6

124 1 7

124 1 8

124 1 9

124 1 10

124 1 12

Entry 129

step 4 should make a OH- not OH radical also its missing a radical arrow from the CH bond to the new OH bond step 5 should have the radical arrow go from the oxygen atom to the iron atom to reset it back to +2 also its missing a radical arrow from the C radical to the new CO bond step 6 in order to make the correct product an extra full arrow is added from a S=O bond to the oxygen

129 1 4

129 1 5

129 1 6

Entry 134

step 1 is missing a radical arrow from O=O to an oxygen atom and the arrow from iron should go to one of the oxygens step 2 and 3 needed to be fused to prevent a 4 Val -1 Nitrogen at the end of step 2's chain of arrows what likely happens is that the arrows travel back towards the oxygens creating the OH an O-2 atom and a Fe+4 atom

134 1 1

134 1 2

134 1 3

Entry 136

the oxygen atom is not shown to be protonated twice in the second step to better have this make sense with the reaction happening around it I split the protonation’s into the first two steps assumed OH3+ was the acid

136 1 1

136 1 2

136 1 3

136 1 4

Entry 137

step 2 has 2 hydrogens on the +1 oxygen this is fixed other steps just needed arrows tweaked

137 1 2

137 1 3

137 1 4

137 1 5

Entry 139

step 1 breaks the Mo S bond and gives both e- to the Mo this results in a -1 charge on the Mo not a -2 this charge difference needed to be updated in step 2 and 3, step 2 is missing the OH- attack on the Mo causing the charge to change by -1 also step 3 should have the arrow come from the Mo O bond instead of the oxygen atom causing a +2 charge change on the Mo resetting it

139 1 2

139 1 3

Entry 140

cobalt’s charge is not supposed to change after homolytic cleavage in step 1 also missing a radical arrow from the CoC bond to the Co, steps 2-9 are updated to have the correct charge of -2, step 4 creates a bond with the Cys H and the nucleotides C2 atom when it should be forming one with the oxygen, step 7 and 8 are missing radical arrows from the SH bond and CH bond respectively, to the new bond

140 1 1

140 1 2

140 1 3

140 1 4

140 1 5

140 1 6

140 1 7

140 1 8

140 1 9

Entry 143

the thioredoxin acceptor is shown is two different ways here I swap out what it looks like in step 5 for what it looks like in step 4

143 1 5

Entry 144

step 1 makes changes to the Mo charge by -1 not -2 so every later step is changed to fit this also step 4 should not change the charge of the Mo only step 5 should add charge back to the Mo to reset it again like in entry 121 I am curious if a Mo+4 atom is ever experimentally observed if so then this mechanism is wrong

144 1 2

144 1 3

144 1 4

Entry 145

step 1 iron gains an e- from the S H bond lowering its charge to +1 and the caption says that water is eliminated but the reaction show produces OH3+ this taken with the HOFe cofactor left at the end suggests that the original complex should be HOFe which would form water if the OFe bond is broken and forms the an OH bond which in thar case would not change the charge on the iron for step 1 I assume the original charge is still +2 homolytic bond breaking or cleavage in steps 2, 3, 6, and 7 do not change the charge of the iron, step 4 should just be a double arrow from iron to Fe O bond causing the iron to lose an e- and gain the charge to make it +3, step 5 should have the iron gain an e- from the Sulfur lowering its charge to +2

145 1 1

145 1 3

145 1 5

145 1 6

145 1 7

Entry 156

actually mech 2 on website, step 1 is a neutral change to the charge of the Ni there is a half arrow missing coming from the S C bond and going to the Ni S bond, step 3 needed Ni charge to be 0, also two half arrows are missing from the Ni S bond going to the S S bond and the Ni atom this still is a neutral change in charge because the product is made in step 3 and the cofactor is reset step 4 is redundant

156 1 1

156 1 2

156 1 3

156 1 4

Entry 159

step 1 should protonate the Asp at the oxygen rather than the carbon, steps 4, 5, and 6 are redundant as the site is already reset after step 3

159 1 1

159 1 4

159 1 5

159 1 6

Entry 195

the amine in the reactant is deprotonated in step 7 but not in the later steps this is changed in steps 1-6

195 1 1

195 1 2

195 1 3

195 1 4

195 1 5

195 1 6

Entry 212

this was the hardest mechanism to fix there were many errors in the databases files the full edited mechanism can be seen in the Marvin files the big change was adding in the discharging of an iron-sulfur cluster each time an ATP is used contiguous steps and charge balance guided the editing

212 1 1

212 1 2

212 1 3

212 1 4

212 1 5

212 1 6

212 1 7

212 1 8

212 1 9

212 1 10

212 1 11

212 1 12

212 1 13

Entry 231

step 2 should use O=O instead of [O][O] and the double bond is homolytically broken with one e- going to the Mn O bond and the other going to the other oxygen this does not change the charge of Mn and the next steps that use Mn need to be updated in order to pull electrons from the carbon in step 5 I had Mn lose an e- in step 4 rather than step 5 step 6 does not change the charge of Mn and the resetting of O=O is missing this is fixed

231 1 2

231 1 3

231 1 4

231 1 5

231 1 6

Entry 239

actually mech 2 on website, step 2 has the OOH os bonded to Fe this is removed, step 3 should has the water grab the proton from Tyr and bond it to the oxygen not to the hydrogen

239 1 2

239 1 3

Entry 250

step 2 and 3 perform the wrong charge changes and the arrows are not going to the right places in balancing the mech I found a simple version that just requires the iron to change from +3 to +2 then back to +3 in steps 2 and 3 rather than the much larger change from +3 to +2 to +1 to -1 that never get reset in the database, step 2 homolytically forms a bond with the iron and oxygen anion while breaking the peroxide bond, step 3 then has the Cl attack the oxygen radical causing the radical to move to the iron changing the charge to +2 step 4 breaks the OFe bond heterolytically protonating the oxygen

250 1 2

250 1 3

250 1 4

Entry 254

actually mech 2 on website methyl group is missing on the cation in step 5 also a hydrogen should be abstracted from that methyl group creating a CC double bond

254 1 5

Entry 261

first step's Marvin file will not open not sure why replaced it with the first step in the 2nd mechanism since it is identical

261 1 1

Entry 264

missing methyl group in step 6 this is fixed in 6 and 7 but the mechanism does not put the double bond on the right carbons as in the overall reaction

264 1 6

264 1 7

Entry 268

this is the second instance of a 3 center 2 e- bond (first was 127) this implies one e- half bonds which cannot be represented as SMILES to fix this I make it a slightly simpler polar mechanism if the Co His bond is covalent then this mechanism can be simplified to have the Co atom and His act as a bridge giving two e- to a water in step 1 and then giving those two e- back to the nitrogen in 5-methyltetrahydrofolate in step 2, lastly the methylated 5-methyltetrahydrofolate is protonated by OH3+

268 1 1

268 1 2

268 1 3

Entry 275

step 7 should have the lone pair on the nitrogen attack the CN bond pushing the CC double bond e- to the oxygen forcing the radical the leave to FAD allowing it to protonate water according to doi:10.1111/j.1742-4658.2009.06966.x the e- is abstracted before the hydrate is formed, step 8 the OH- ion attacks the keto carbon realizing the acetate

275 1 7

275 1 8

Entry 276

Mo should not lose charge in step 1 this effects every later step the catalyst is not reset at the end but in balancing the mechanism by modifying the charge on the Mo using the proper arrows reveals that the charge is reset on the Mo this oddly predicts that the Mo goes through two steps where it is +7 charge

276 1 2

276 1 3

276 1 4

276 1 5

276 1 6

276 1 7

Entry 278

cytochrome protonation and radicalization are not shown since step 2 is swaps an e- between equally charged Cu atoms there is no overall change I instead completed the cytochrome protonation assumed OH3+ was the proton donors

278 1 1

278 1 2

Entry 281

steps 1 and 5 are missing radical arrows in sink FAD makes more sense for step 3 to make a carbon cation in the FADH then the transformation in step 6 makes sense but because it is a radical in step 6 the product file made is invalid and will not open, so this fix was done manually

281 1 1

281 1 3

281 1 5

281 1 6

Entry 297

The Asp should stay deprotonated in the first step

297 1 1

Entry 323

step 2 has to create a N Fe bond homolytically which in combination with the e- coming from the iron sulfur cluster changes the irons charge by -1 this is updated for every heme intermediate in the later steps, step 5, 8, and 11 do not have a lone pair to grab to allow for the nitrogen to be protonated the original file gets around this by putting an alias showing N+ when the nitrogen really is neutral instead I put the protonation in the next step also just so ammonium is made I protonated ammonia in step 13

323 1 3

323 1 4

323 1 5

323 1 6

323 1 7

323 1 8

323 1 9

323 1 10

323 1 11

323 1 12

323 1 13

Entry 358

iron sulfur cluster did not change in the 1 step resulting in the state being wrong in step 10 also the radical arrows for step 1 and 10 are wrong it should ba a single e- is donated by the cluster to the methyl radical group then the C S bond donate its two e- to the sulfur letting it be neutral PLP cofactor was still not reset stills needs a possible protonation step but I am not sure where

358 1 1

358 1 4

358 1 5

358 1 7

358 1 10

Entry 370

step 1, 4, and 5 should not change the charge on the iron, step 2 adds 1 charge and step 3 subtracts 1 charge also made the mechanism demethylate the single methylated lysine as this is the reactant in the overall reaction

370 1 2

370 1 3

370 1 4

370 1 5

370 1 6

Entry 374

in order to reset catalyst step 6 should make a O=V bond with the V+ O- bond to minimize charge getting it back to its original valence, step 7 should protonate the other oxygen anion

374 1 6

374 1 7

Entry 409

first step is missing a curved arrow where the C=O bond forms a OH bond using the amine H step 6 does not need an external acid to protonate the nitrogen a H is missing on the alcohol

409 1 1

409 1 6

Entry 538

step 3 deleted a proton on the amine

538 1 3

Entry 545

Pyridoxal 5'-phosphate nitrogen in the ring should be protonated in steps 1 and 2

545 1 1

545 1 2

Entry 550

step 1 is redundant as the Lys and Asp end in deprotonated and protonated states, step 2 missing a curved arrow where the C=O bond forms a OH bond using the amine H

550 1 1

550 1 2

Entry 551

this step forgot to protonate the C4a-hydroxyflavin this assumes it was by a OH3+ ion so that and water are cancelled out

551 1 3

Entry 553

step 2 forgot to keep the Lys protonated and have it protonate the ketone while it attacks C2, step 3 assumed that the ketone was protonated twice in step 2 instead it should be protonated once then protonated again in step 3, step 6 missed arrows swapping a proton from the water to the Lys

553 1 2

553 1 3

553 1 6

Entry 562

step 2 only adds one e- to the Mo due to the Mo O bond breaking, one e- is lost in step 3 also an iron sulfur cluster is replaced as the e- sink because an R group cannot be generalized to work with other rules, step 4 should not change the charge on the Mo due to it forming a bond and losing an e-

562 1 3

562 1 4

Entry 563

step 1 the formate is assumed to bond to the Mo changing it to a +5 state, step 2 the formate is deprotonated by a His forming CO2 breaking the Mo O bond changing the Mo state to +4, step 4,5,and 6's Marvin files would not open so they had to be edited assumed step 4 uses a OH3+ ion to protonate the menaquinone

563 1 1

563 1 2

563 1 3

563 1 4

563 1 5

563 1 6

Entry 578

missing the formation of bonds due to the removal of metallic bonds here they are just shown as bonds formed from polar chemistry

578 1 1

578 1 2

578 1 3

578 1 4

Entry 600

the irons do not change its charge at any point in this mechanism there is only homolytic bond breaking and forming the iron oxygen complex is not reset there needs to be two added steps where a NADH donates its H- to the oxygen while both of the O Fe bonds are broken homolytically forming a OH- then the OH- is protonated

600 1 3

600 1 4

600 1 5

600 1 6

600 1 7

600 1 8

600 1 9

Entry 613

added Asp Lys proton swap step for step 2 and pushed the later steps forward

613 1 2

613 1 3

613 1 4

613 1 5

613 1 6

613 1 7

Entry 618

step 1 forgot have the O=C grab a H from the amine

618 1 1

Entry 657

step 2 shows a full arrow going from a coordination bond this is removed

657 1 2

Entry 660

the iron atom does not gain its charge in the first step instead step 3 is when the Fe O bond is broken causing it to lose an e- this assumes that the Glu14 does not bond to the Fe in step 3, it appears to have only a binding bond in the products

660 1 2

660 1 3

Entry 683

the single paper referenced has the mechanism start with the C3 alcohol deprotonated DOI:10.1021/bi015652u

this allows it to form the reactant in step 2

683 1 1

Entry 685

step 1 should start with a O=O homolytically breaking and forming a FeO bond homolytically, step 2 should have the real bond between O2 and iron and have a homolytic bond formation to the keto group of 2-oxoglutarate but this does not change the charge on iron, step 3 should form the CO2 bond from the lone pair not the binding connection, step 4 and step 5 should have a homolytic bond breaking of O Fe which does not change the charge of the iron, step 6 should change the charge of iron by -1 due to the lone pair of Oxygen donating the e- to the bond since this happens an added step is needed to reset the irons charge and unbinds the OH and succinate

685 1 1

685 1 2

685 1 3

685 1 4

685 1 5

685 1 6

685 1 7

685 1 8

Entry 694

NADH electron donation step is not shown NADH first donates an Hydride to the FAD making FADH- then this FADH- donates its Hydride to FMN making FMNH- these two steps are what step 1 and step 2 are in the new mechanism, step 3 takes one e- from the FMNH- to bind oxygen to the heme iron creating FMNH radical and Fe+2, step 4 takes the other e- from FMNH radical to fill the lone pair on the oxygen, step 5 and 6 reduce the oxygen and resets the heme iron to +3, step 7 protonates the substrates alcohol group, step 8 abstracts the H on the substrate, step 9 homolytically breaks the FeO bond forming the alcohol group on the substrate

694 1 1

694 1 2

694 1 3

694 1 4

694 1 5

694 1 6

694 1 7

694 1 8

694 1 9

Entry 699

NADH electron donation step is incorrect, NADH first donates an Hydride to the FAD making FADH-, then this FADH- donates its Hydride to FMN making FMNH-, these two steps are what step 1 and step 2 are in the new mechanism, step 3 takes one e- from the FMNH- to bind oxygen to the heme iron creating FMNH radical and Fe+2, step 4 takes the other e- from FMNH radical to fill the lone pair on the oxygen, step 5 and 6 reduce the oxygen and resets the heme iron to +3,step 7 abstracts the H on the substrate, step 8 homolytically breaks the FeO bond forming the alcohol group on the substrate

699 1 1

699 1 2

699 1 3

699 1 4

699 1 5

699 1 6

699 1 7

699 1 8

Entry 702

step 4 and 5 are missing a hydrogen on the nitrogen in ring

702 1 4

702 1 5

Entry 709

the radical arrows are shown wrong for step 2 the iron donated one e- to the new double bond with another from C=C bond and then the oxygen donated a lone pair cause the iron to gain an e- lowering its charge step 3 should be where the iron atom donates an e- so as to cancel out the cation on the Trp allowing for the radicals on the heme and the Trp to be reformed into a double bond in step 4 the same goes for step 5 the iron atom donates an e- so step 7 can reform the double bond step 6 protonates the oxygen by breaking the FeO bond resetting the heme iron

709 1 2

709 1 3

709 1 4

709 1 5

709 1 6

709 1 7

Entry 711

Cys needs to be deprotonated before attacking

711 1 1

Entry 712

thiolate should be deprotonated before attacking in both steps

712 1 1

712 1 2

Entry 713

nitrogen’s pass a hydrogen in step 1 and 5 the oxygen should not start protonated in step 3 but should be protonated by an acid in this case I assume its OH3+

713 1 1

713 1 3

713 1 5

Entry 722

step 3 forgets to protonate the oxyanion I assume it is done by a OH3+ ion, step 5 forgets to show that the adjacent alcohol protonated the new oxyanion, step 6 is missing a double headed arrow

722 1 3

722 1 5

722 1 6

Entry 730

nitrogen’s pass a hydrogen in step 1 and 9

730 1 1

730 1 9

Entry 737

Lys should be protonated in step 1, a protonation should happen before step 5 so since step 4 is empty I put the protonation of the intermediate there, the Lys should be protonated and the homolytic bond breaking should not have changed the Co charge in step 1 and it does not change during homolytic bond forming in step 8, the Lys should also be protonated in steps 8 and 9

737 1 1

737 1 4

737 1 7

737 1 8

737 1 9

Entry 742

missing a step where the PLP cofactor is reset this added step returns the last bond to the PLP to the Lys and has the phosphate protonate the amine on the product also step 2 has an OH3+ donate a proton to the Lys to make sure it is at the correct protonation state

742 1 2

742 1 11

Entry 743

step 2 should use O=O and homolytically breaks to form the Cu O bond but this does not change the charge on the copper every step after needed the charge corrected on the copper this also resets copper

743 1 2

743 1 3

743 1 4

743 1 5

743 1 6

Entry 745

missing a His deprotonation in step 6

745 1 6

Entry 753

step 1 and 2 miss the oxygen protonation from the amine and that the nitrogen connected to the ring should start protonated as in steps 3, step 3 misses the nitrogen protonation from the oxygen, step 4 should have the oxygen protonate the amine to get it in the carbonyl state its in for step 5

753 1 1

753 1 2

753 1 3

753 1 4

Entry 758

step 1 does not have the correct radical arrows there should be a homolytic OO bond breaking with one of the e- going to the new O Fe bond this should not change the charge on the Fe, step 2 should have the radical arrow go from the double bond to the new OC bond, step 3 has Fe gaining an e- losing a charge but in step 4 the Fe O bond should have broken releasing water and having Fe gain a charge resetting

758 1 1

758 1 2

758 1 3

758 1 5

Entry 759

PLP cofactor should have its oxygen protonated initially in steps 2 and 3

759 1 2

759 1 3

Entry 762

nitrogen in ring should be protonated in step 2 and the Lys should not be protonated in step 8 so as to reset the PLP

762 1 2

762 1 8

Entry 768

step 1 arrow missing for the water, step 5 the nitrogen should get protonated by His175

768 1 1

768 1 5

Entry 792

Cys should be deprotonated before it attacks

792 1 1

Entry 797

there has to be some type of base that takes a proton from the lys I assume its water

797 1 1

Entry 809

forgets the protonation of the nitrogen by the water in steps 4 and 5

809 1 4

809 1 5

Entry 855

Cys reactant should have a protonated carboxyl in steps 1, 2, and 3, PLP oxygen should not be protonated in step 3 as it is in the rest of the mechanism

855 1 1

855 1 2

855 1 3

Entry 857

the carbonic acid should be deprotonated by water not itself in step 2 in order to make the reactant in the next step

857 1 2

Entry 867

to better fit with other rules the general halogen is replaced with Cl

867 1 1

Entry 894

for some reason the mechanism is not shown in the database at the time of writing this 8-3-25 balancing the steps implies that the reactant is trimethylammonia not trimethylammonium meaning the overall reaction is backwards from what the mechanism shows this lack of a protonation is needed for the nitrogen to attack to bond with one of the oxygens on the Mo and breaking its bond then changing the charge on Mo by -1, the next step breaks the Mo bond severing it from Mo making the product trimethylamine N-oxide and changing the charge on Mo by +1 but a oxygen is not added to the Mo resetting it this is likely done by water attacking and being deprotonated and 2 Fe+3 accepting the 2 extra e- two extra steps were added to reset this cofactor

894 1 1

894 1 2

894 1 3

894 1 4

Entry 907

the carbamylated lysine should be deprotonated by water in step 1 and 2 if it is to be consistent with the later steps, step 4 is only consistent with the reactants of step 5 if a OH- attacks instead of water

907 1 1

907 1 2

907 1 4

Entry 918

actually mech 2 on website the Tys are likely to exchange their protons for the radical

918 1 1

Entry 925

the Mo does not lose two charges in step 1 this effects every later step and properly resets the Mo charge

925 1 1

925 1 2

925 1 3

Entry 943

step 1 takes an e- from the iron-sulfur cluster making that atom (+) and the S-adenosyl radical then that S-adenosyl radical is give back in step 10 resetting the cluster, steps 1, 2, 3, 4, and 5 are missing radical arrows

943 1 1

943 1 2

943 1 3

943 1 4

943 1 5

943 1 10

Entry 949

the iron sulfur cluster donates an e- to the SAM neutralizing the +1 charge of the S and creating the radical everything else in the mech seems fine but I added the first two steps in reverse as the last two steps this way the SAM iron sulfur cluster is reset this way the overall reaction is just the demethylation of the methylphosphonate this also balances the overall reaction before it was off by 2 e-

949 1 1

949 1 5

949 1 7

949 1 8

Entry 968

the charge on Mo does not change in step 3 or step 4, step 5 does have a charge change to reset the cofactor OH- attacks an extra step where the lone pair on the FADH finishes the creation of FADH2

968 1 4

968 1 5

968 1 6

Entry 981

iron sulfur clusters do not change in step 2 causing them to be wrong in later steps also His protonation step between step 3 and 4 is missing this is fixed by adding a OH3+ protonation to step 4 but this cancels out the His for the step

981 1 3

981 1 4

Entry 986

heme is not used in this mechanism instead I put two full heme molecules in the mechanism, step 2 donates each e- to two Fe+3 ions, step 3 creates the product tetrathionate

986 1 1

986 1 2

986 1 3

Entry 987

Mo should not change charge in step 2 or 4, step 4 makes more sense if the full arrow from the S H bond goes to the Mo S bond rather than the Mo atom since the Mo=S bond is already created step 5 should only add charge to the Mo then step 6 has water donate its lone pair to the Mo O lowering the change and resetting the cofactor

987 1 4

987 1 5

987 1 6

Entry 995

actually mech 3 on website, step 1 should not change the charge of the Mn so all later steps that use Mn needed to be changed, step 2 has the Mn losing a charge as in the database but step 6 should have it gain back the charge when the Mn O bond breaks

995 1 3

995 1 5

995 1 6

S1.3 Labeling codes for catalytic groups

To distinguish between reaction substrates/products and reusable catalytic groups (such as amino acid side chains and cofactors), we employ a system of labeling codes. These codes identify specific moieties that are exempt from the cumulative sum constraint within the optimization formulation, reflecting their dynamic and recyclable role as part of the enzyme's machinery. The list below details all the labeled moieties used in this study. Those with ‘none’ next to them are given a label but are never catalytically used in the M-CSA database.

1 water/oxonium/hydroxide

-Canonical amino acids

2 His

3 Asp/Glu

4 Lys

5 Cys

6 Tyr

7 Ser

8 Arg

9 Asn/Gln

10 Thr

11 Gly

12 Phe

13 Trp

14 Met

15 Ala

16 Pro

17 Leu 'none'

18 Val 'none'

19 Ile 'none'

-Non-canonical amino acids

20 Trq tryptophan tryptophylquinone residue

21 Cso cysteine-hydroxide residue

22 Sec selenocysteine residue

23 Kcx carbamylated lysine residue

24 Pyr pyruvoyl residue

25 Fgl formylglycine residue

26 Sep phosphorylated serine residue 'none'

27 Hyp proline with an alcohol group residue 'none'

28 Csd cysteine-sulphinic acid residue 'none'

-Metals

29 Mg Magnesium

30 V Vanadium

31 Mn Manganese

32 Fe Iron

33 Co Cobalt

34 Ni Nickel

35 Cu Copper

36 As Arsenic

37 Se Selenium

38 Mo Molybdenum

39 Hg Mercury

-Cofactors

40 FAD/FMN

41 NAD(P)(H)

42 Pyridoxal 5-phosphate

43 Heme

44 Iron-Sulfur cluster

45 Thiamine Diphosphate

46 Molybdopterin

47 Glutathione

48 Vanadate

49 SAM

50 Cobalamin

51 Biotinate

52 Pyrroloquinoline Quinone

53 Dipyrromethane cofactor

54 Iminodimethanethiolate

55 Acetic Acid

56 Coenzyme F430

57 Dioxygen

58 2-[(1S)-1-aminoethyl]-1-carboxymethyl-5-hydroxy-4-methylimidazole

S2. Neural Network Re-ranking Analysis

As stated in the main text, we first attempted to use a deep neural network to re-rank the top-ten parsimonious mechanisms. This section provides the details of that analysis.

S2.1. Methods - Neural Network Architecture

The network was designed to take a matrix representation of a candidate mechanism as input, where rows represent unique moieties and columns represent the elementary rules in that mechanism (Figure S2). A series of dense layers create a learned feature embedding for each rule, which are then summed in a permutation-invariant aggregation step. This single vector representation is then processed by a final classifier to produce a probability score between 0 and 1.

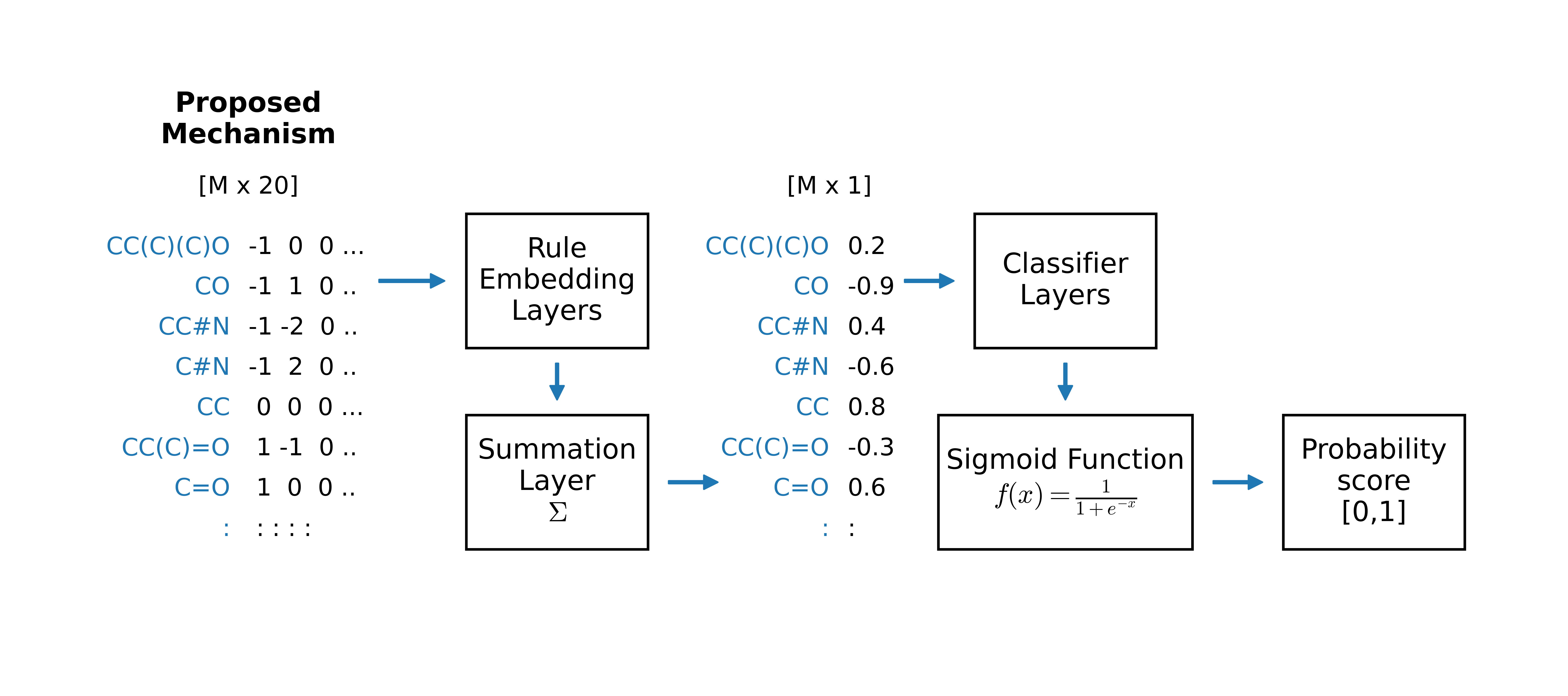

Figure S2. Architecture of the tested neural network re-ranker. The model processes a matrix representation of a candidate mechanism to produce a single probability score used for re-ranking.

S2.2. Results and Discussion

The model was trained and evaluated using a 5-fold cross-validation scheme on the M-CSA dataset. The results (Figure S3) show that while the network learned to distinguish some chemical features, it failed to provide an overall improvement in the recovery of the correct mechanism at rank 1 compared to the parsimony-only baseline. This result led us to develop the alternative similarity-based re-ranking method described in the main text.

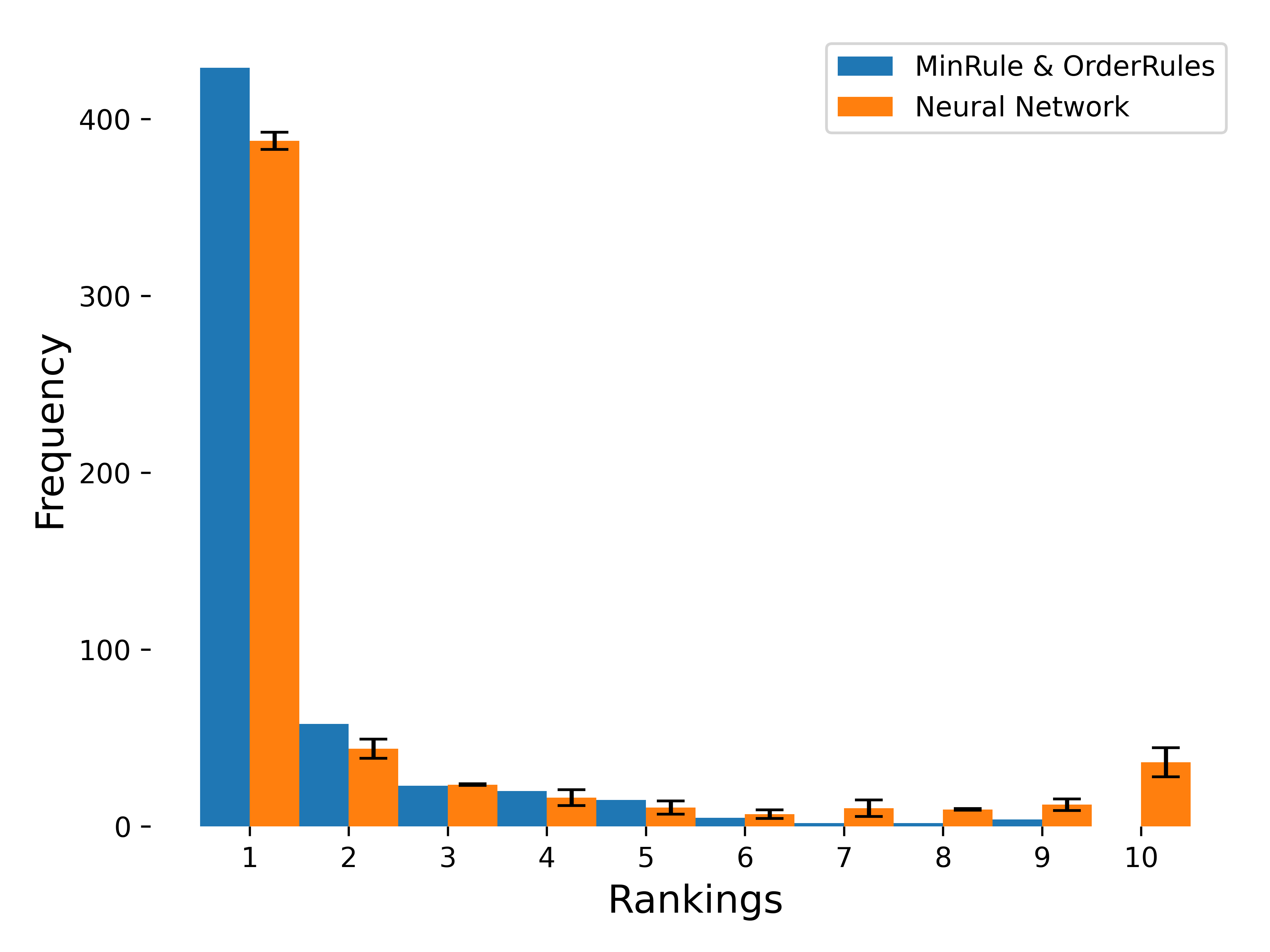

Figure S3. Performance comparison of the parsimony-based framework and the neural network re-ranker. The plot shows the distribution of ranks for the top ten predictions across the 661 M-CSA test cases. The orange bars (Neural Network) do not show a significant improvement over the blue bars (parsimony-only), particularly at rank 1.
